# Identification of Novel Inhibitors of JEV RdRp as Potent Antiviral Drugs: Targeting NS5-NS3 Protein Interaction

**DOI:** 10.64898/2026.09.23.753689

**Authors:** Preeti Mishra, Shriyanshi Mishra, Gaurava Srivastava, Km. Archana, Mohammad Imran Siddiqi, Sourav Haldar, Raj Kamal Tripathi

## Abstract

Japanese Encephalitis Virus (JEV) belongs to the Flavivirus family, and the RNA-dependent RNA polymerase (RdRp) domain located at the C-terminus of non-structural protein 5 (NS5) regulates de novo viral genome synthesis. The conserved priming loop inserted in the thumb domain of RDRP initiates de novo genome synthesis. Interestingly, the reported replication process initiates with an interaction between NS5 (methyltransferase/RDRP) and NS3 (protease/helicase), and inhibiting this interaction directly correlates with the inhibition of viral replication. The function of the priming loop in the context of the NS5-NS3 interaction is not yet known. In this study, we studied the priming loop function in the NS5-NS3 interaction and viral replication. Using a structure-based drug design approach, we screened the Maybridge compound library against the RdRp priming loop and identified 15 candidate compounds based on binding energy. Among them, DSHS00151 showed significant dose-dependent inhibition of the NS5-NS3 interaction with an IC50 of 3.5 µM in the mammalian two-hybrid assay, inhibited viral infectivity with an IC50 of 2.54 µM, and reduced viral RNA load with an IC50 of 2.9 µM, compared to CD10712. Molecular Dynamics (MD) simulations revealed that DSHS00151 exhibits stronger and more stable binding with the priming loop residues (790–812) compared to CD10712. The molecular mechanism suggested by the docking of the NS5-NS3 proteins showed that the priming loop tends to rotate toward the NS3 protein to stabilize the complex, and compound binding restricts this loop rotation, thus compromising the stability of the NS5-NS3 complex and affecting viral replication. Overall, our study validated the function of the NS5 priming loop as an allosteric site, and compounds binding to this allosteric site belong to the Non-Nucleoside Reverse Transcriptase Inhibitors (NNRTI) class of inhibitors, which disrupt the NS5-NS3 interaction and could be developed as novel anti-JEV therapeutics.

## 1. Introduction

Japanese encephalitis is caused by the Japanese encephalitis virus, an arbovirus in the genus Flavivirus. This causes severe infection of the central nervous system and is responsible for the major outbreak of vaccine-preventable encephalitis in many parts of Asia and the Western Pacific. The first JE case ever reported was in Japan in 1871(Quan et al., 2020) Outbreaks have been occurring every 10 years. In 1924, the very first outbreak of JEV that occurred recorded more than 6,000 cases and 3,000 reported deaths in 6 weeks. In Asia, several outbreaks of JE cases were recorded subsequently(Erlanger et al., 2009) . Recently, in 2005, in the northern part of India and Nepal, large outbreaks were reported with 5000 cases and took a toll on 1300 lives and in the current scenario, 24 endemic countries are found to be at high risk of JE infection, risking 3 billion lives (Quan et al., 2020). Approximately 11 kb genome of JEV contains only one open reading frame encoding a single polyprotein, which is subsequently cleaved by cellular and viral proteases to yield three structural proteins (SPs) and seven non-structural proteins (NSPs) (NS1, NS2a, NS2b, NS3, NS4a, NS4b, NS5). Upon entering the host cell, JEV releases its genomic RNA into the cytoplasm, where it undergoes initial translation using host cell machinery to produce a polyprotein, which is cleaved by various host and viral proteases; this is followed by replication of the JEV genome. The SPs are incorporated into virus particles, whereas the NSPs participate in the formation of the replication complex (RC), assembly complex (AC) and the replication of the viral genomic RNA(S. Kumar et al., 2022).NS5 protein is constituted by two distinct domains as well, namely an N-terminal methyltransferase and a C-terminal RNA-dependent RNA polymerase that are required for capping and synthesis of the viral RNA genome, respectively (Wang et al., 2009).NS3 and NS5 proteins are the major enzymatic components of the viral replication complex, which promotes efficient viral replication in close association with cellular host factors. Several studies have suggested that NS5 and NS3 exhibit enzymatic activity and physically interact within the replication complex, contributing to viral replication. (Brand et al., 2024) (Osawa et al., 2023). Due to their numerous functions and their central role in the virus life cycle, NS3 and NS5 have been designated as important drug targets(Sampath & Padmanabhan, 2009). The synthesis of the JEV RNA genome occurs de novo, meaning that it is a primer-independent viral replication process, which is a common feature of the Flavivirus family. Flaviviruses possess a unique priming loop located within the thumb domain of the RNA-dependent RNA polymerase (RdRp)(Lu & Gong, 2013). The priming loop, which spans amino acid residues 790 to 812, an extension of the thumb subdomain that projects towards the palm-domain active site, plays a crucial role in stabilising the closed pre-initiation conformation(Lu & Gong, 2013). It also provides a platform for the synthesis of new RNA strands. As the process transitions to elongation, the priming loop and the surrounding fingers, along with the thumb and palm channels, undergo conformational changes that allow for RNA extension. Furthermore, the interface between the palm and thumb forms an allosteric site that can regulate the conformational change between initiation and elongation(Godoy et al., 2017). However, the stabilization of the priming loop during replication initiation is not well understood. It has been suggested that the interaction between NS3 and NS5 proteins plays a crucial role in this process(Brand et al., 2017). NS3, known for its helicase and protease activities, may be thought to interact with NS5, the RNA-dependent RNA polymerase, to stabilize the priming loop. This interaction may enhance the structural integrity of the RNA replication complex, facilitating a smoother transition from initiation to elongation. Additionally, the binding of NS3 to the priming loop may prevent premature dissociation of the RNA template, allowing for efficient synthesis and ensuring proper replication fidelity. Understanding the dynamics of the NS3-NS5 interaction could, therefore, provide insights into the regulatory mechanisms that govern the stability of the priming loop during viral replication.

In this study, we developed a mammalian two-hybrid assay to quantify the interactions between the NS3 and NS5 proteins, in which the involvement of the conserved priming loop in the interaction could be analyzed (Figure 1D). We docked 55,717 Maybridge compounds onto the priming loop region and selected the top 15 compounds based on their binding energies (Table 1). Next, we evaluated the inhibition of NS5-NS3 protein interactions in the presence of these 15 selected compounds (using a single concentration 12.5µM) using the mammalian two-hybrid assay (Table 2). We identified two compounds, DSHS00151 and CD10712, that exhibited differential dose-dependent inhibition of NS3-NS5 interactions in the mammalian two-hybrid assay (Figure 2). Furthermore, we examined the effects of DSHS00151 and CD10712 on viral replication and viral load and found dose-dependent reductions in infectivity and replication. (Figure 3). Molecular Dynamics (MD) analysis revealed that DSHS00151 has a higher binding affinity to the priming loop compared to CD10712 (Figure 5). The molecular mechanism proposed by Docking studies of the NS5-NS3 indicated that the NS5 RDRP priming loop rotates to engage with the NS3 protein, which is crucial for stabilization (Figure 8). DSHS00151 binds to the priming loop with high affinity, restricting the priming loop from rotating to bind with NS3, ultimately leading to an unstable replication complex. Overall, our study characterized a novel allosteric site on the RNA-dependent RNA polymerase (RdRp) protein, and inhibitors targeting the priming loop may represent a new class of non-nucleoside reverse transcriptase inhibitors (NNRTIs) that could be developed as novel anti-JEV therapeutics.

**Figure 1:**
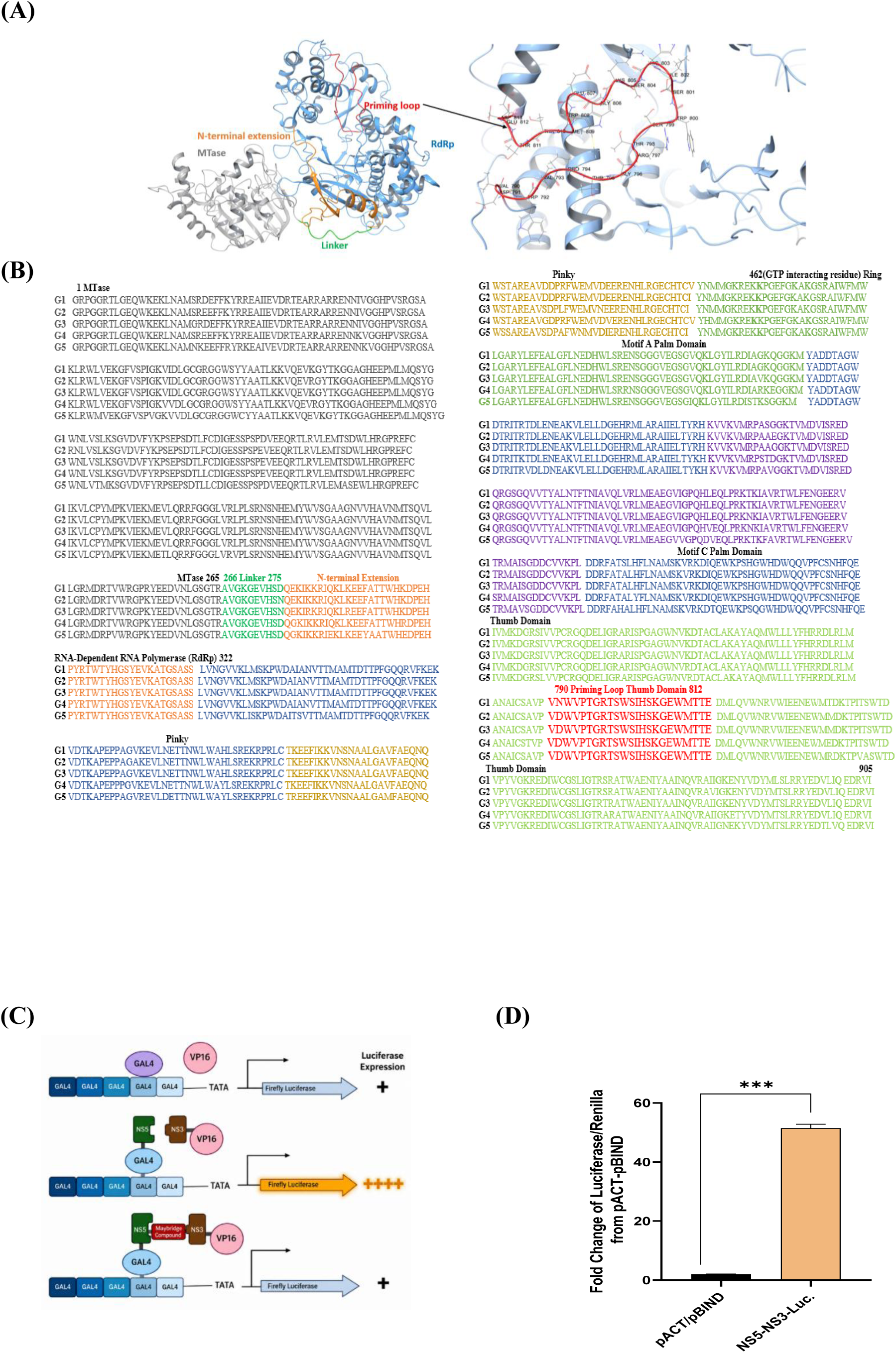
(A) The crystal structure of full-length Japanese Encephalitis Virus (JEV) NS5 (PDB 4K6M) is shown, viewed from above the RNA-dependent RNA polymerase (RdRp) active site, with a colour-coded bar indicating the various structural elements. The colouring scheme is as follows: MTase (grey), the linker (green), N-terminal extension (orange), priming loop (red), and RdRp (blue). (B) The protein sequences from five geographical isolates (Japan G1, Australia G2, US G3, Indonesia G4, and China G5) were aligned to identify conserved and variable residues. The positions of the MTase(grey), linker (green), RdRp, and their respective sub-domains (e.g., Palm (olive green), Thumb (Light green), Priming Loop (Bold Red), Pinky (Blue), Ring) are indicated above the sequence blocks. This alignment reveals a high degree of sequence conservation across the catalytic core of the RdRp domain. (C) **Basal Reporter System (Lane 1):** Five GAL4 binding sites upstream of a TATA box drive the expression of a Firefly Luciferase reporter gene. Unbound promoter sites or non-interacting GAL4 DNA-binding domains yield basal-level expression (**+**). **Direct Interaction Assembly (Lane 2):** GAL4 DNA-binding domain fused to NS5 recruits the NS3-VP16 activation domain fusion via direct NS5–NS3 complex formation, driving high luciferase reporter gene expression (**++++**). **Competitive Inhibition Assay (Lane 3):** Addition of a small molecule (Maybridge Compound) disrupts the interaction between the NS5 and NS3 proteins, preventing VP16 recruitment to the promoter and reducing reporter gene expression back to basal levels (**+**). (D) The interaction between NS5 and NS3 was evaluated using a mammalian two-hybrid assay. HEK293T cells were co-transfected with mammalian two-hybrid constructs encoding the NS5 and NS3 proteins, along with a luciferase reporter plasmid. Control groups included pBIND-pACT empty vectors. The interaction between NS5 and NS3 (NS5–NS3-LUC) resulted in an increase in reporter gene expression compared to the controls, indicating a specific protein–protein interaction. Data are presented as a mammalian two-hybrid ratio (relative luciferase activity) and represent the mean ± SEM from independent experiments.

**Figure 2.**
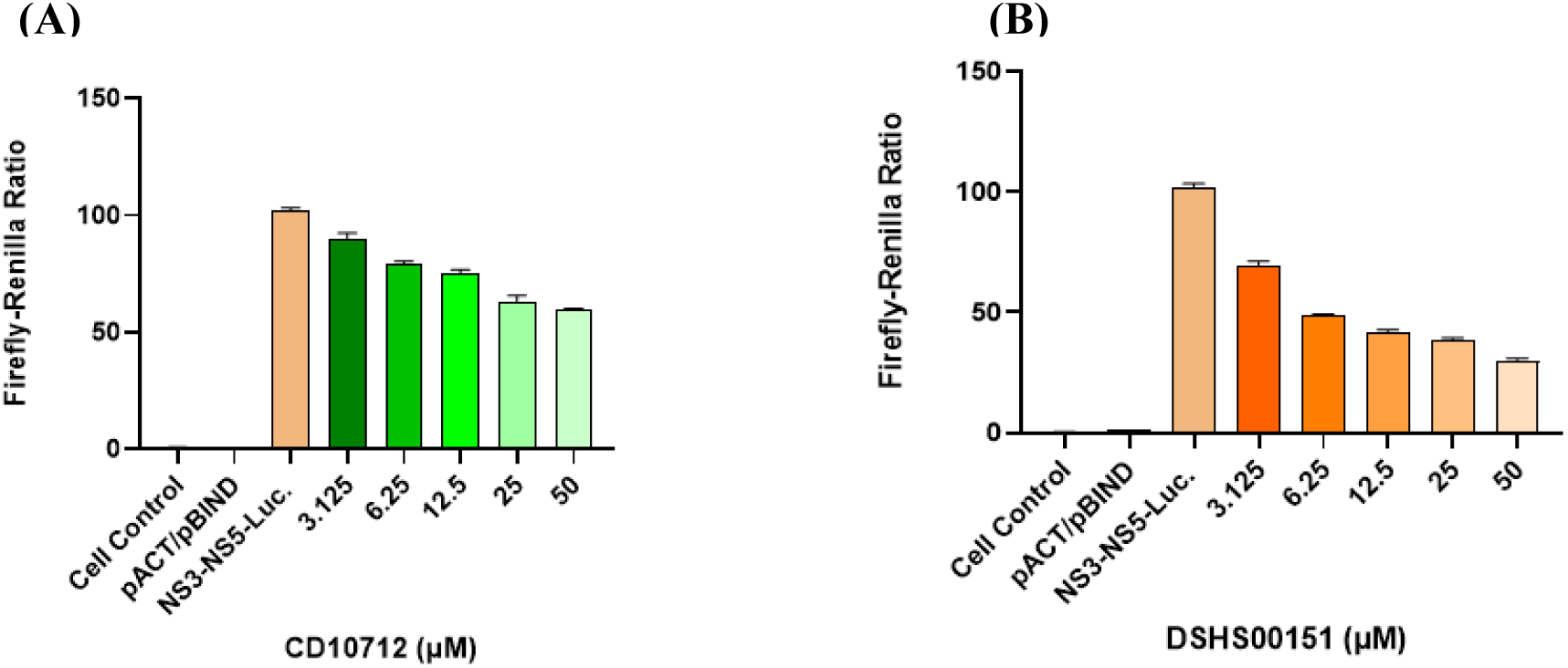
Dose-dependent inhibition of NS5-NS3 interaction by CD10712 and DSHS00151 in a mammalian two-hybrid assay. (A) The effect of increasing concentrations of CD10712 (3.125–50 µM) on the interaction between NS5 and NS3 proteins was assessed. The reporter activity decreased progressively as the concentration of the compound increased, demonstrating a dose-dependent inhibition of the NS5–NS3 interaction. (B) The effect of increasing concentrations of DSHS00151 (3.125–50 µM) on the NS5–NS3 protein interaction was evaluated. Cells were co-transfected with the specified constructs, and the strength of the interaction was quantified using dual luciferase reporter activity (Mammalian Two-Hybrid Ratio). A concentration-dependent reduction in reporter activity was observed, indicating a disruption of the NS5–NS3 interaction. Data are presented as Firefly - Renilla ratio (relative luciferase activity) and represent ± SEM from independent experiments.

**Figure 3.**
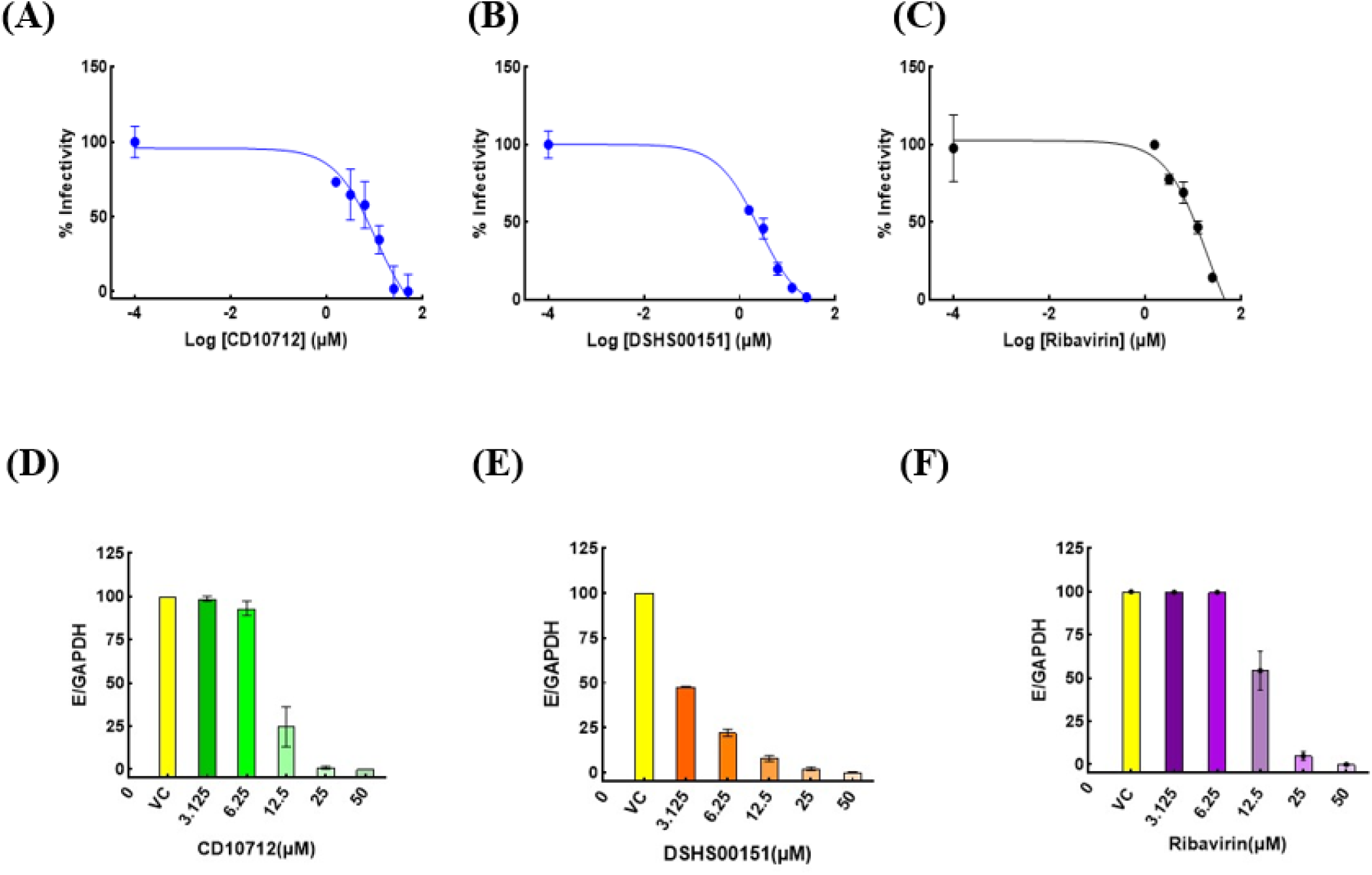
Effect of CD10712 and DSHS00151 on JEV infection. (A) Flow cytometry analysis of CD10712, DSHS00151, and Ribavirin (used as positive control) on JEV infection. Data represent the mean ± SEM from at least three independent experiments. (B) CD10712, DSHS00151, and Ribavirin show estimation of viral RNA content in the presence of test compounds using qRT-PCR. The virus-only group represents infected, untreated cells. The viral Envelope (E) gene was normalized using GAPDH as an internal control. Data presented ± SEM from two independent experiments.

**Table 1:**
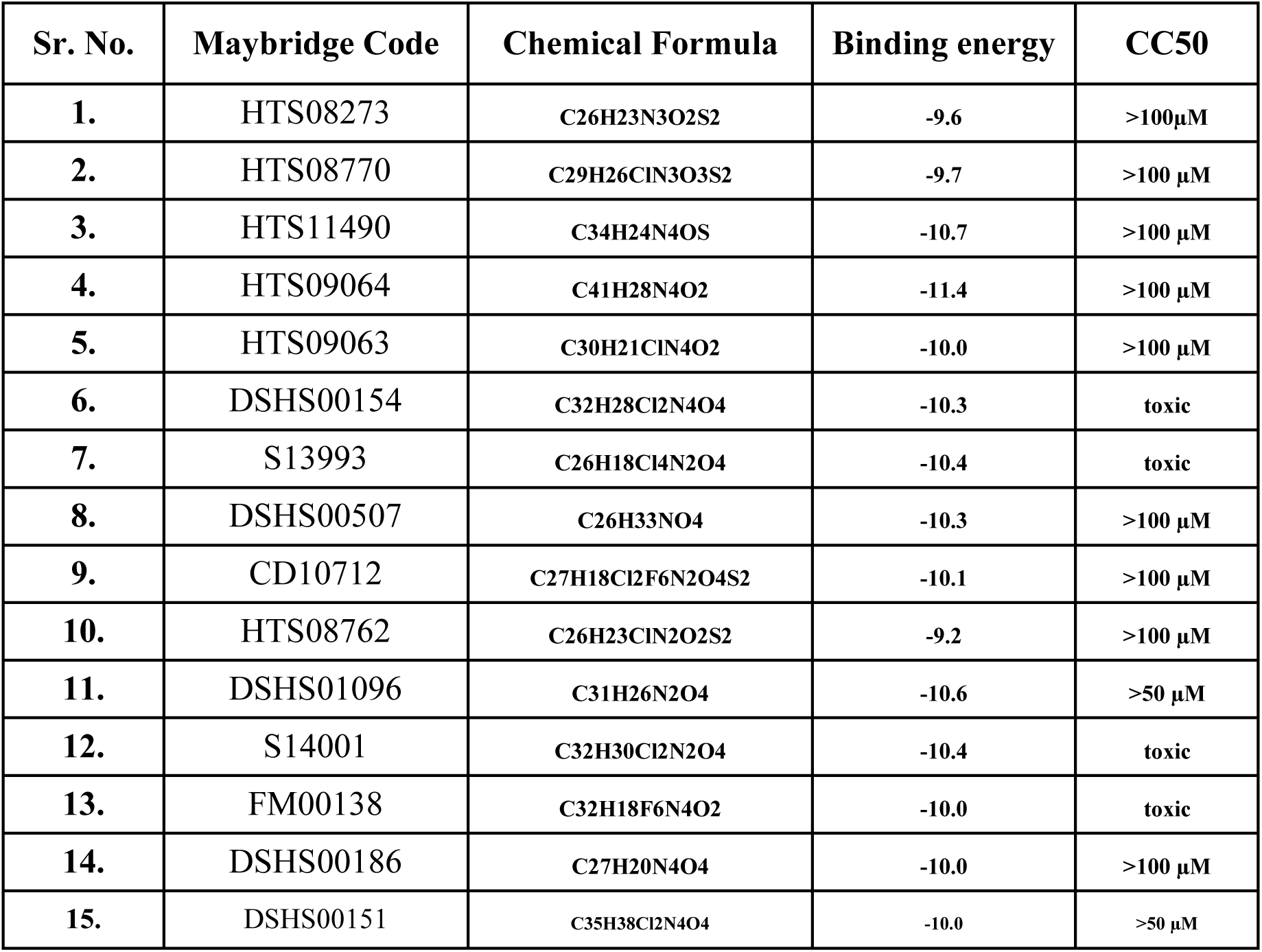
List of top 15 Maybridge compounds based on binding energy at the JEV RdRp priming loop, analyzed for cytotoxicity (CC_50_)

**Table 2.** Mammalian Two-Hybrid Evaluation of NS5–NS3 Interaction Inhibition by Compounds.

| S.No. | Compound Name | Firefly-Renilla Ratio | Fold Change =<br>Compound Ratio /<br>pACT-pBIND Ratio | Fold reduction<br>in luciferase<br>expression<br>compared to<br>untreated cells |
| --- | --- | --- | --- | --- |
| 1. | pACT-pBIND-Luc. | 2.06 | - | - |
| 2. | NS5-NS3-Luc. | 101.90 | 49 ± 0.85 | 1.00 |
| 3. | HTS08273 | 99.53 | 47.82 ± 0.46 | 1.02 |
| 4. | HTS09064 | 87.5 | 41.53 ± 0.91 | 1.17 |
| 5. | S13993 | 109.21 | 53.04 ± 0.07 | 0.92 |
| 6. | HTS08762 | 101.75 | 48.58 ± 0.77 | 1.00 |
| 7. | S14001 | 103.28 | 50.36 ± 0.26 | 0.98 |
| 8. | DSHS00186 | 93.33 | 44.78 ± 0.49 | 1.09 |
| 9. | HTS08770 | 89.41 | 42.39 ± 0.98 | 1.14 |
| 10. | HTS09063 | 75.26 | 35.93 ± 0.58 | 1.32 |
| 11. | DSHS00507 | 105.41 | 50.19 ± 0.93 | 0.96 |
| 12. | DSHS01096 | 101.82 | 49.46 ± 0.08 | 1.00 |
| 13. | FM00138 | 97.44 | 47.02 ± 0.24 | 1.04 |
| 14. | CD10712 | 75.19 | 36.01 ± 0.46 | 1.36 |
| 15. | HTS11490 | 88.83 | 42.13 ± 0.95 | 1.14 |
| 16. | DSHS00154 | 90.83 | 43.30 ± 0.75 | 1.11 |
| 17. | DSHS00151 | 43.51 | 20.84 ± 0.26 | 2.33 |

Quantification of NS5–NS3 protein-protein interaction inhibition and relative luciferase expression fold change in response to compound treatment. The empty vector control (pACT-pBIND-Luc) represents basal transcriptional activity, whereas the untreated NS5-NS3-Luc complex serves as the positive interaction shows (49) fold higher compared to control. Compounds CD10712 and DSHS00151 differentially disrupt the NS5–NS3 interaction, reducing interaction levels to 1.36 and 2.33 folds compared to positive interaction control respectively. Data are presented as Firefly– Renilla ratio (relative luciferase activity) and represent ± SEM from independent experiments.

## 2. Materials and methods

### 2.1 Protein sequence alignment of JEV NS5 genotypes

Based on the NCBI Reference Sequences: JAPAN (G1) AAF73859.1, AUS (G2) NP_059434.1, US (G3) BAE48356.1, INDO (G4) AAP39942.1, and CHINA (G5) AEK75355.1, we performed a multiple sequence alignment using Molecular Evolutionary Genetics Analysis (MEGA) software. The protein domains are represented as follows: MTase (grey), the linker (green), N-terminal extension (orange), and RdRp (blue). The priming loop region is highlighted in bold red (Figure 1B).

### 2.2 Cell culture and cytotoxicity assay

Vero Cells (Vero Cells C1008, cat. CRL-1586, ATCC) were cultured in growth medium [DMEM (D5648-L, Sigma) with 10 % FBS (10270-106, GIBCO)] under 5 % CO2 at 37 ◦C.

Cells were seeded in a 96-well cell culture plate (1×10^4^ cells/well) in 200 μl/well. After 12 h of seeding, Compounds were added at different concentrations in triplicate with a vehicle (DMSO) control. The plate was incubated at 37◦C with 5 % CO2. At 68 h, MTT (3-(4,5-Dimethylthiazol-2-yl)-2,5-Diphenyltetrazolium Bromide, Sigma) was added. The plate was incubated at 37 ◦C with 5 % CO2. At 72 h, the supernatant was removed carefully, and DMSO (100 μl/well) was added to dissolve the formazan crystals. The plate was kept under shaking conditions for 10–15 min. Absorbance was measured at 570 nm using a microplate reader (Varioskan LUX, Thermo Scientific)

### 2.3 RNA extraction from JEV-infected and uninfected cells

Vero cells were cultured and seeded in 6-well plates at a density of 0.3 × 10⁶ cells per well in Dulbecco’s Modified Eagle Medium (DMEM) supplemented with 10% Fetal bovine serum (FBS) and incubated at 37°C in a humidified environment with 5% CO₂. After 24 hours, cell seeding, the cells were infected with Japanese encephalitis virus (JEV; Vellore Strain) at a multiplicity of infection (MOI) of 0.01. Following a 2-hour viral adsorption period, the inoculum was discarded, and fresh DMEM supplemented with 2% FBS was added. The infected cells were incubated for 72 hours. The cells were harvested and lysed for total viral RNA isolation using standard TRIzol reagent (Thermo Fisher Scientific).

### 2.4 Mammalian two-hybrid assay

Protein-protein interaction between JEV NS3 and NS5 was examined using the Check Mate™ Mammalian Two-Hybrid System (Promega). HEK293T cells were seeded 24 h before transfection. Cells were co-transfected with pBIND-NS5, pACT-NS3, and the pG5luc reporter plasmid using TurboFect Transfection Reagent (Thermo Fisher Scientific, Waltham, MA, USA). Plasmids were transfected at a 1:1:1 ratio, with 750 ng of each plasmid per transfection. DNA-TurboFect complexes were prepared according to the manufacturer’s instructions, incubated for 20–30 min at room temperature, and added dropwise to cells maintained in complete 1X DMEM supplemented with 10% fetal bovine serum.

The experimental design included untransfected cells as the cell control, empty pACT and pBIND vectors as the negative control, and the MyoD and ID plasmid pair supplied with the kit as the positive control.

Following transfection, cells were incubated for 48 h. Cells were then lysed using 1× Passive Lysis Buffer (Promega), and luciferase activity was measured using the Dual-Luciferase® Reporter Assay System (Promega). Firefly luciferase activity was first measured following addition of Luciferase Assay Reagent II, after which Stop & Glo® Reagent was added to quench firefly luciferase activity and measure Renilla luciferase activity. Luminescence was recorded using a GloMax® luminometer (Promega). Firefly luciferase activity was normalised to Renilla luciferase activity, and the normalised luciferase ratio was used to evaluate the interaction between NS5 and NS3.

#### 2.4.1 Preparation of cDNA and cloning of JEV NS5 and NS3 genes in mammalian two-hybrid vectors

First-strand cDNA synthesis was performed from the total RNA extracted from JEV-infected Vero cells using a commercial cDNA synthesis kit (G2P). Instead of kit-supplied oligo(dT) or random primers, cDNA synthesis was done with a combination of the gene-specific NS5 wild-type forward primer (5’-TAGGATCCGGAAGGCCTGGCGGAAGGAC-3’, engineered with a BamHI restriction site) and the reverse primer (5’TAGAATTCTCAAATGACCCTATCCTCCTGAATCAGCACG-3’, engineered with an XbaI restriction site). The synthesized cDNA was subsequently used as a template for polymerase chain reaction (PCR) amplification employing the same set of NS5 wild-type forward and reverse primers. The amplified PCR product was resolved on a 1% agarose gel, excised under UV transilluminator, and purified using the G2P Cutting Edge Gel Extraction Kit. The purified NS5 wild-type amplicon and the destination pBind vector backbone (CheckMate™ Mammalian Two-Hybrid System, Promega) were digested with BamHI and XbaI restriction enzymes, gel-purified, and ligated using T4 DNA Ligase to generate the recombinant pBind-NS5 wild-type construct.

Similarly, total RNA extracted from JEV-infected Vero cells was used for cDNA synthesis using commercial cDNA synthesis kit (G2P). The NS3 gene was amplified using gene-specific primers comprising the NS3 forward primer (5’-GAGGATCCGGGGCGTGTTTTGGGACAC-3’, containing a BamHI site) and NS3 reverse primer (5’-TCTCTAGATCTCTTCCCTGCTGCAAAGT-3’, containing an XbaI site). The NS3 amplicon was resolved by 1% agarose gel electrophoresis, excised, and purified. Both the purified NS3 PCR product and the pAct vector backbone (Promega) were subjected to double restriction digestion with BamHI and XbaI, gel-purified, and ligated using T4 DNA Ligase to yield the recombinant pAct-NS3 expression vector.

#### 2.4.2 PCR amplification and intermediate TA subcloning of NS5

Full-length JEV NS5 plasmid (BIOMATIK) was amplified using the NS5 forward primer (5’-TAGGATCCGGAAGGCCTGGCGGAAGGAC-3’) and reverse primer (5’-TAGAATTCTCAAATGACCCTATCCTCCTGAATCAGCACG-3’). PCR Cycling was performed on a Bio-Rad C1000 Thermal Cycler. Amplicons were resolved on 1% agarose gels, and gel purification was done using the G2P Cutting Edge Gel Extraction Kit. Purified DNA concentrations were quantified using a Varioskan LUX multimode reader (Thermo Scientific). Purified amplicons were inserted into the pGEM®-T/pGEM®-T Easy vector system (Promega). Ligation products were transformed into competent E. coli DH5α cells and plated on LB agar containing kanamycin (50 µg/mL), IPTG, and X-Gal for blue/white screening. Positive colonies were picked and cultured overnight, after which plasmid DNA was isolated using the Spin Clean Plasmid Prep Kit (G2P).

### 2.5 Maybridge compound screening

The virtual screening of the Maybridge Library against JEV RNA-dependent RNA polymerase (RdRp) was performed using the PyRx software integrated with AutoDock Vina. The three-dimensional structure of the protein JEV-RdRp (PDB id. 4K6M) was retrieved and prepared by removing water molecules, ligands and ions, followed by appropriate preprocessing before docking. The Maybridge compound library, comprising 55,717 molecules, was imported into PyRx, where the compounds were energy-minimised using the Universal Force Field (UFF) and converted into PDBQT format. The active site of RdRp was defined based on the known catalytic residues, and the grid box was generated to encompass the catalytic domain. Virtual screening was conducted using AutoDock Vina with appropriate exhaustiveness parameters, and binding affinity (kcal/mol) was recorded for all the molecules. The docking poses with an RMSD value of 0.0 Å were considered for further analysis, and compounds were ranked based on their binding affinities. The top 15 molecules exhibiting the highest binding affinities were selected and tabulated.

#### 2.5.1 Molecular docking

To better understand the molecular mechanism of active compounds, selected active compounds were redocked and taken ahead for MD simulation. Initial preparation of protein and ligands was done through AutoDock Tool 1.4.5. Molecular docking was performed through AutoDock 4.2(Morris et al., 2009). Docking parameters included: maximum number of generations: 27,00,000, Number of evaluations: 2,50,00,000, Population size: 200 and GA runs: 200. The grid box was generated by covering the priming loop region with 0.375 Å of grid spacing. Lamarckian Genetic Algorithm (Huey et al., 1998) was used for docking pose generation.

#### 2.5.2 Molecular dynamics simulation of JEV NS5 (PDB 4K6M)

MD simulation studies were performed for 100ns each with GROMACS 4.6.2 (Hess et al., 2008) (Van Der Spoel et al., 2005) using the CHARMM36 all-atom force field (Huang & Mackerell, 2013). Topologies of ligand ere generated through the SwissParam server with the MM force field approach (Bugnon et al., 2023). Each system was placed in the centre of a cubic box having a distance of 1.0 nm between the protein and the edge of the simulation box, and TIP3P, an explicit water molecule, was used to solvate. All the systems were neutralized by adding Na+ or Cl− ions accordingly and then minimized with the steepest descent method. Two nanoseconds of equilibration were performed at constant volume, pressure (1 atm) and temperature (300K). LINCS method was used to restrain the bond length, and the Particle Mesh Ewald (PME) algorithm (Hess et al.) was used with a grid spacing of 1.6 Å and a cut-off of 10 Å to calculate the electrostatic interaction. Finally, 200ns of production MD was done for each system and trajectories were saved every 20 ps.

#### 2.5.3 Trajectory analysis

Gromacs analysis modules, g_rms, g_rmsf and g_hbond, were used to analyse root mean square deviation (RMSD), root mean square fluctuation (RMSF) and hydrogen bonds. H-bond occupancy was calculated through a Perl script, and total residual contribution (as donor + as acceptor) was used for the result analysis. Plots were generated using Xmgrace. Visual analysis and figure preparation were performed using PyMOL.

#### 2.5.4 Binding energy calculation

Binding free energy estimation of all the docked complexes was done by the MM/PBSA method through the g_mmpbsa module. For each simulated system, 200 snapshots were taken over the last 40 ns of the trajectory at 200 ps intervals. The module calculates electrostatic interactions, van der Waals interactions, polar solvation energy and non-polar solvation energy.

### 2.6 Primary screening of docked compounds using the mammalian two-hybrid assay

Compounds selected through virtual screening and molecular docking were initially evaluated for their ability to disrupt the NS5-NS3 interaction using the mammalian two-hybrid assay. HEK293T cells were seeded one day before transfection and transfected as described above. Two hours after transfection, individual compounds were added at a final concentration of 12.5 µM. Vehicle-treated cells served as controls.

Following 48 h of incubation, cells were lysed with 1X Passive Lysis Buffer, and Dual-Luciferase® Reporter Assays were performed. Relative luciferase activity was determined after normalization of firefly luciferase to Renilla luciferase. Compounds producing a significant reduction in normalized luciferase activity compared with the vehicle control were considered primary hits.

### 2.7 Dose-dependent inhibition of the NS5–NS3 interaction

Compounds identified during the primary screen were further evaluated in dose-response experiments using the mammalian two-hybrid assay. HEK293T cells were transfected as described above, and compounds were added 2 h post-transfection at final concentrations of 50µM-3.125µM. Following 48 h of incubation, cells were lysed, and firefly and Renilla luciferase activities were measured using the Dual-Luciferase® Reporter Assay System.

Relative luciferase activity was calculated as the Firefly/Renilla ratio, and dose-response bar graphs were plotted to evaluate concentration-dependent inhibition of the NS5-NS3 interaction.

### 2.8 Flow cytometry

Vero cells were seeded in a 96-well plate, followed by infection with JE virus [Japanese Encephalitis Virus (Vellore Strain), a kind gift from Dr Vikas Agrawal at the Sanjay Gandhi Postgraduate Institute of Medical Sciences (SGPGIMS), Lucknow] at 0.01 MOI. The plate was incubated at 37°C for 2 h under humid conditions in 5% CO2. At 2 h post-infection, infection media was removed, and fresh media at 200μl/ well (DMEM with 2% FBS) with compounds at different concentrations (25µM-0.78µM) was added. The plate was incubated under humid conditions at 37°C in 5 % CO2 for 72 h. Overlay media was aspirated, and cells were washed with 1x PBS, 150 µl/well. Cells were harvested by adding 0.25% trypsin-EDTA (25 µl/well). Next, cells were transferred to a U-bottom 96-well plate and were fixed with 4 % paraformaldehyde (PFA) (50 µl/well). After proper mixing, the plate was incubated for 30 min at room temperature (RT) to fix the cells. Cells were permeabilised with 1x permeabilisation buffer at 150 µl/well (2% BSA + 0.1% saponin in PBS). Next, 40 µl/well blocking solution (1 % normal rabbit serum in 1x permeabilisation buffer) was added. After 30 min of incubation at RT, 4G2-Alexa488 mAb (4G2 antibody against the envelope fusion domain of Flavivirus, Novus company) (20 µl/well) was added, and the antibody dilution was prepared in blocking buffer (1:1500). The Plate was incubated for 1 h at 37 ◦C with shaking. After that, the cells were washed with 1X permeabilisation buffer and pelleted by centrifugation at 1000g. The cell pellet was resuspended in 1x PBS (200 µl/well). The samples were read on a flow cytometer (BD Bioscience FACS LYRIC). Data were analysed using FlowJo software (BD Bioscience).

### 2.9 Quantification of viral RNA by RT-qPCR

Vero cells were seeded (0.3 × 10^6^ cells/well) in a 6-well culture plate. Cells were infected with JEV at 0.01 MOI. At 2 h post-infection, the various concentrations of compounds (50μM-3.125μM) were added. After 72 h of infection, cells were collected. Total RNA extraction was carried out using the Monarch Spin RNA Isolation Kit (New England Biolabs). cDNA synthesis was performed with the Verso cDNA Synthesis Kit (Thermo Fisher Scientific AB-1453/A) using the C1000 Touch Thermal Cycler (Bio-Rad). 0.5 μg of total RNA was used for cDNA synthesis. RT-qPCR was performed in the Real-Time GreenR PCR Master Mix 2X (G2P) using Bio-Rad universal Syber Green Mastermix (1725120) and 500 ng of cDNA. For RT-qPCR, primers were designed and purchased from Integrated DNA Technologies, Inc. (IDT) for the structural gene, Envelope (E) of JEV; E(FP)-5 ′ TGCATGGAACCACCACTT3 ′; E(RP)-5 ′ GAAGGAGCATTGGGTGTTATTG 3’ (97bp).

Statistical analysis of the results normalised to GAPDH was carried out using one-way ANOVA, followed by Dunnett’s multiple comparisons test using GraphPad Prism.

## 3. Results

### 3.1 Elucidation of the JEV priming loop region in the RdRp protein and development of mammalian two-hybrid assay quantitating NS5-NS3 interaction in HEK-293T cells

The JEV NS5 protein crystal structure shows a natural fusion protein with the priming loop highlighted in red (Figure 1A) (Lu & Gong, 2013). All JEV genotypes’ NS5 proteins circulate in different geographical regions, were aligned and compared, revealing that all NS5 proteins were highly conserved (Figure 1B). The JEV NS5 protein is composed of methyltransferase residues 1 to 265, RdRp residues 322 to 905, and linker residues 266-275. All RdRp motifs were highly conserved and distinguished by different colours (Figure 1B). The unique residues in the RdRp priming loop, specifically positions 790-812, are conserved across all JEV genotypes. These residues are located in the thumb region, which extends into the active site. This extension is sometimes supported by a C-terminal extension of the thumb, facilitating the RdRps in the de novo synthesis of RNA (Lu & Gong, 2013).

In JEV-infected cells, the NS5 protein has been shown to interact with the NS3 protein, forming a replication complex that regulates viral RNA replication (Brand et al., 2017). The NS5 and NS3 residues and domains that are involved in the interaction were characterized, and mutations of interacting residues showed inhibition of interaction, resulting in inhibition of viral replication. (Tay et al., 2015; Nannetti et al., 2025).The RdRp domains of NS5 must be aligned to interact with the NS3 protein to form a functional replication complex (Brand et al., 2017;Osawa et al., 2023). The functional role of the conserved priming loop in the NS5-NS3 replication complex in regulating viral replication is not well known. To understand it, we first developed an NS5-NS3 interaction assay mimicking the replication complex in virus-infected HEK-293T cells, using a mammalian two-hybrid system in which NS5-NS3 physical interaction was quantified by luciferase expression (Figure 1C & D). The mammalian two-hybrid assay has been extensively used to identify interacting protein domains during interactions and cellular function regulated by the interaction (B. Kumar et al., 2013; P. Kumar et al., 2019).

The NS5 gene was cloned into the pBind, and the NS3 gene was cloned into the pAct mammalian two-hybrid vectors. The physical interaction of NS5 and NS3 will complete the transcription circuit and express the luciferase reporter vector in cells. The inhibitors inhibiting the interaction could not express the luciferase reporter (Mendonça et al., 2013). The luciferase gene expression demonstrates the strength of physical interactions among expressed proteins within cells (Figure 1D). In our study, to demonstrate NS5-NS3 interaction, we co-transfected HEK-293T cells with the NS5 pBIND vector and the NS3 pAct vector along with a luciferase reporter vector. As a control, we co-transfected HEK-293T cells with empty pAct and pBind vectors along with the luciferase reporter vector. Our results showed that the HEK-293T cells co-transfected with NS5 and NS3 vectors showed ∼49-fold luciferase/renilla expression compared to cells transfected with the pAct/pBind control (Figure 1D). Overall, our findings demonstrate that the JEV NS5-NS3 interaction has been developed in HEK-293T cells, which resembles the replication complex in infected cells, and could be effectively used to characterize the RdRp domains involved in interaction with the NS3 protein.

### 3.2 In silico screening of the Maybridge compound library against the RdRp priming loop and cytotoxicity evaluation

In our previous experiment (Figure 1D), we developed the NS5-NS3 interaction complex in HEK-293T cells using a mammalian two-hybrid system. To understand the function of the priming loop during the NS5-NS3 interaction and JEV replication, we needed pharmacological inhibitors that could effectively inactivate the priming loop activity. We used a structure-based drug development approach to screen the Maybridge compound library against the RdRp priming loop (residues 790-812). The virtual screening of the Maybridge Library against the JEV-RdRp priming loop was performed using the PyRx software integrated with AutoDock Vina. The three-dimensional structure of the protein JEV-RdRp (PDB id. 4K6M) was retrieved and prepared by removing water molecules, ligands and ions, followed by appropriate preprocessing before docking. The Maybridge compound library, comprising 55,717 molecules, was imported into PyRx, where the compounds were energy-minimised using the Universal Force Field (UFF) and converted into PDBQT format. The active site of RdRp was defined based on the known catalytic residues, and the grid box was generated to encompass the catalytic domain. Virtual screening was conducted using AutoDock Vina with appropriate exhaustiveness parameters, and binding affinity (kcal/mol) was recorded for all the molecules. The top 15 molecules exhibiting the highest binding affinities were selected and tabulated (Table 1). Additionally, these compounds were analyzed for cytotoxicity. All compounds were tested at concentrations of (100µM-1.56 µM) on Vero cells. The CC50 results for the compounds are presented in Table 1.

### 3.3 Analysis of Maybridge compounds for inhibition of NS5–NS3 interaction and antiviral activity

The small molecules screened against the RdRp priming loop were selected based on their binding energy (Table 1). To understand that these compounds may have the potential to inhibit the NS5-NS3 interaction, thereby impacting viral replication. We screened 15 compounds to evaluate their ability to disrupt the NS5-NS3 interaction and their antiviral activity. In our experiment, we used a single non-toxic concentration of 12.5 µM to assess the effects of the compounds on the NS5-NS3 interaction. The NS5pBind and NS3pAct vectors were co-transfected along with a luciferase reporter vector. Control setups included co-transfecting the pAct and pBind empty vectors with the luciferase reporter in HEK-293T cells. All co-transfected cells containing the NS5 and NS3 vectors were treated with or without the compounds at a concentration of 12.5 µM, and luciferase expression was quantified thereafter. Our results showed (Table 2) that the co-transfected cells containing NS5 and NS3 without treatment exhibited a luciferase expression level of ∼49-fold compared to the control with the pAct and pBind vectors, whereas the effect of the compounds on NS5-NS3 interaction, and interaction dependent luciferase expression is shown in Table 2.

Specifically, upon comparing the fold luciferase expression in cells treated with CD10712 and DSHS00151 to untreated cells, it was reduced by 1.35 and 2.34 times, respectively (Table2). The differential reduction in luciferase expression suggests that these compounds may bind the priming loop with different affinities, leading to varying degrees of NS5-NS3 interaction inhibition.

Subsequently, all 15 compounds were screened for their anti-JEV activity. We treated JEV-infected cells with the compounds at the same non-toxic concentration of 12.5 µM for 72 hours. The results were summarized in Table 3. When comparing treated to untreated cells, CD10712 and DSHS00151 showed 0.33 and 0.17 times reduction in virus infectivity, respectively (Table3).

**Table 3.** Screening of Candidate Small-Molecule Compounds for anti-JEV Activity.

| S.No. | Compound Name | Infectivity (percentage) | Fold reduction of infectivity compared to untreated cells |
| --- | --- | --- | --- |
| 1. | Virus Control | 60.25 ± 4.75 | - |
| 2. | HTS08273 | 64.30 ± 1.20 | 1.05 |
| 3. | HTS09064 | 59.40 ± 1.00 | 0.93 |
| 4. | S13993 | 69.20 ± 1.50 | 1.13 |
| 5. | HTS08762 | 65.25 ± 0.65 | 1.06 |
| 6. | S14001 | 63.30 ± 1.20 | 0.98 |
| 7. | DSHS00186 | 78.75 ± 1.15 | 1.28 |
| 8. | HTS08770 | 61.15 ± 1.05 | 0.96 |
| 9. | HTS09063 | 43.70 ± 1.80 | 0.66 |
| 10. | DSHS00507 | 62.90 ± 1.40 | 0.98 |
| 11. | DSHS01096 | 61.55 ± 1.55 | 1.01 |
| 12. | FM00138 | 63.90 ± 1.00 | 1.03 |
| 13. | CD10712 | 21.95 ± 1.05 | 0.33 |
| 14. | HTS11490 | 65.45 ± 0.85 | 1.06 |
| 15. | DSHS00154 | 55.25 ± 2.25 | 0.91 |
| 16. | DSHS00151 | 11.40 ± 1.10 | 0.17 |

Percent viral infectivity in JEV-infected cells following treatment with identified compounds at a fixed screening concentration (12.5 μM). Untreated viral control serves as the baseline for infectivity (60.25%). Compounds CD10712 and DSHS00151 demonstrate the most pronounced reduction in viral infectivity, lowering infection rates to 20% and 10%, respectively. The fold change of virus infectivity upon CD10712 and DSHS00151 compound treatment is 0.33 and 0.17 respectively compared with virus control. Data are presented as infectivity (percentage) and represent ± SEM from independent experiments.

In summary, our findings indicate that a prime loop in the RdRp protein influenced by these two compounds, leading to reduced NS5-NS3 interaction resulting inhibition of virus replication.

### 3.4 Dose-response effects of CD10712 and DSHS00151 on NS5–NS3 interaction

In our previous experiment, we identified two compounds, CD10712 and DSHS00151, that exhibited differential effects on NS5-NS3 interaction, resulting in differential antiviral activity. We analyzed the dose-dependent interaction-inhibition response of DSHS00151 and CD10712 to demonstrate the direct relationship between the intensity of NS5-NS3 interaction and antiviral activity. In this experiment, we evaluated the dose-dependent effect of the compounds CD10712 and DSHS00151 at concentrations (50µM-3.125µM) (Figure 2A & B). pBindNS5 and pActNS3 were co-transfected with a luciferase reporter plasmid into HEK-293T cells, which were then treated or untreated with the compounds at different concentrations. After 48 hours post-transfection, the cells were harvested, lysed, and luciferase expression was quantified.

Our results indicated that both compounds, CD10712 (Figure 2A) and DSHS00151 (Figure 2B), demonstrated a dose-dependent reduction in luciferase expression in NS5-NS3 co-transfected cells compared to the control. The values analysed in Graph pad prism 8.0.2 identified the interaction inhibition concentration (IC50) calculated for CD10712 was 8.701 µM, and for DSHS00151 it was 3.5µM. This suggests that the efficacy of DSHS00151 for interaction inhibition is significantly higher compared to that of CD10712. Overall, these results confirm that both CD10712 and DSHS00151 interfere differentially with the priming loop, resulting in differential NS5-NS3 interaction, demonstrating RdRp priming loop maintains NS5’s ability to interact with NS3.

### 3.5 Dose-dependent effects on antiviral activity and viral RNA load

To better understand the relationship between NS5-NS3 interactions responsible for viral replication, we analyzed the effect of two compounds, CD10712 and DSHS00151, at concentrations (50 µM to 3.125), on virus infectivity and viral replication, using ribavirin, an RdRp inhibitor, as a positive control. JEV-infected cells were treated with the compounds at various concentrations for 72 hours. Viral infectivity was assessed using a flow cytometry assay, while viral RNA replication was evaluated through qRT-PCR.

Our results showed that the CD10712 compound had an IC50 of 10.45 µM (Figure 3A), whereas the DSHS00151 compound demonstrated an IC50 of 2.54 µM (Figure 3B). In comparison, ribavirin, the positive control known to inhibit RdRp, exhibited an IC50 of 19.98 µM (Figure 3C). The ribavirin, known RdRP inhibitor used as positive control both in vitro and invivo studies was used as positive control in our study (Sebastian, et al, International Journal of Antimicrobial Agents 33 (2009) 168–173). These findings suggest that DSHS00151 showed significantly reduced viral infectivity compared to CD10712.

Furthermore, to substantiate whether this reduction in viral infectivity stemmed from direct suppression of viral RNA synthesis, we evaluated the effect of both compounds, CD10712 and DSHS00151, at concentrations of 50µM to 3.125 µM on viral RNA synthesis using qRT-PCR. Dose-response curves generated by plotting relative fold change against compound concentration revealed that DSHS00151 inhibited viral RNA replication with an IC50 2.9µM (Figure 3E) and CD10712 with an IC50 18.32µM (Figure 3D).

Overall, our results establish a strong relationship based on dose-dependent response. The differential binding of compounds at the priming loop inhibits viral infectivity and viral replication.

### 3.6 Computational characterization of CD10712 and DSHS00151 interactions with the priming loop

Initial RMSD trajectories (Figure 4) of the protein backbone have shown a stable trajectory throughout the 100ns. Although DSHS00151 exhibited a small shift around 50–60 ns, it remained stable throughout the rest of the simulation time. If we compare the RMSF trajectories of all the systems, fluctuations in multiple regions around the AAs 270, 440, 570, 601 and 791-810 were found to be restricted in comparison to the RdRp (Apo) system and were aligned with their IC50 trend. Out of these 5 regions, the regions around AA 601 and 791-810 were only lying within the active site of RdRp and may have direct contact with the inhibitors, whereas restricted movement around the rest of the regions was not a direct impact of ligand binding and may have been observed because of the synergistic effect of ligand binding. If we focus on the region around AA 601 and 790-810, both regions supported ligand binding from both ends. At the same time, it was also observed that both regions, around AA 601 and 791-810, were highly stable in the RdRp + DSHS00151 system, representing a strong binding of compound DSHS00151 with an arrested motion of the loop from AA 791-810. In contrast, regions around AA 601 and 799-810 were found with comparatively high fluctuations in the RdRp + CD10712 system. Such fluctuations at both ends of compound CD10712 may be observed due to less stable contact between either or both ends and compound CD10712. This finding supports our previous experimental results, where we have found an increased IC50 value for compound CD10712 in comparison to compound DSHS00151. Further H-bond trajectories have also depicted that the RdRp + DSHS00151 system has a better hydrogen bond pattern in comparison to the RdRp + CD10712 system. Although both systems have only 1-2 hydrogen bonds, the bonding pattern was found to be better and more continuous in the RdRp + DSHS00151 system.

**Figure 4.**
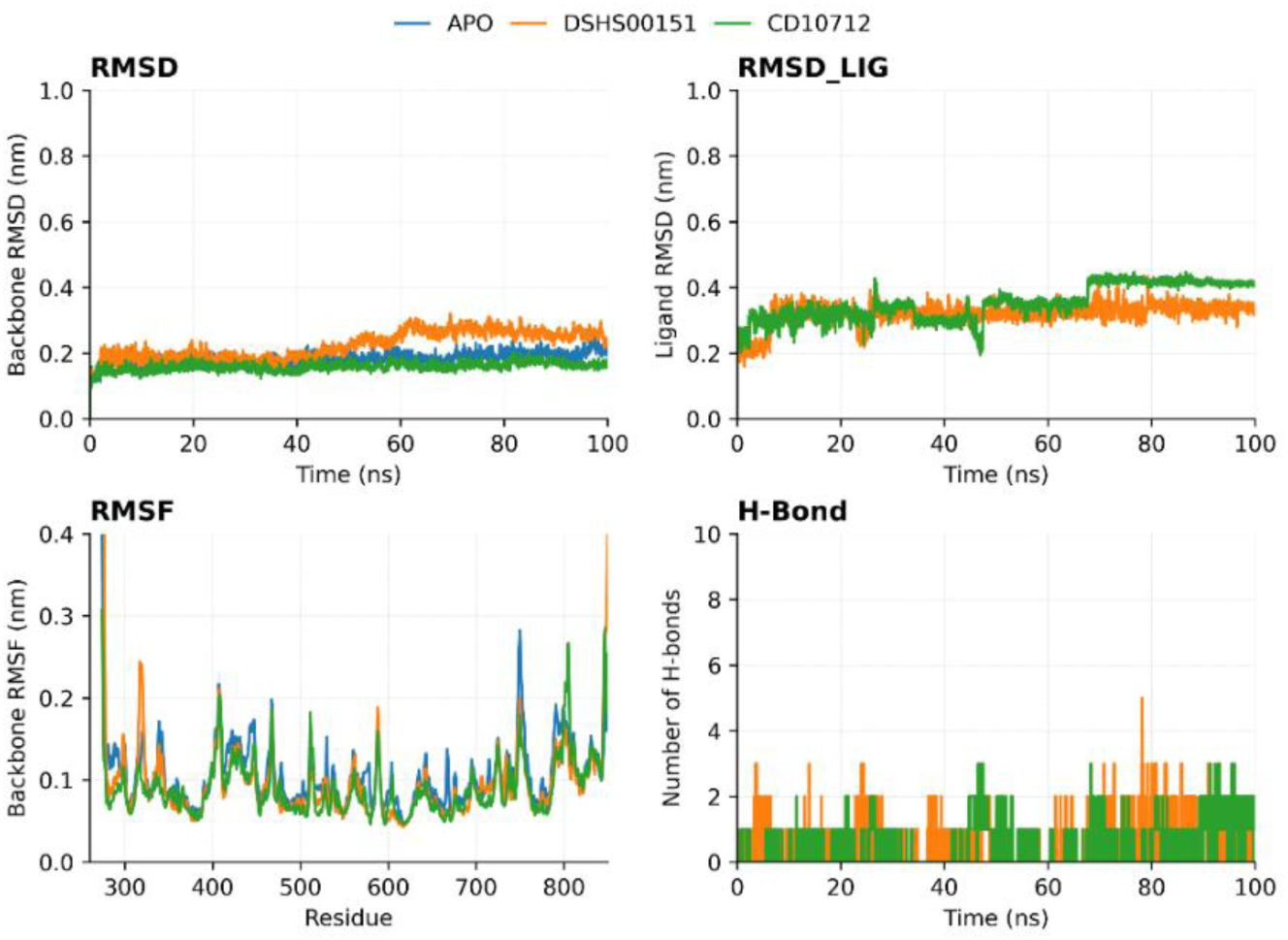
Different analyses for 100ns of MD trajectories; systems are shown in different colours, as mentioned in the legend. (For interpretation of the references to colour in this figure legend, the reader is referred to the web version of this article.)

Thereafter, the contribution of residues was investigated through H-bond occupancy results (Figure 5A), which showed that residue SER-801, which belongs to the same characterised loop discussed before, was a highly contributing residue for H-bond formation in the RdRp + DSHS00151 system, but no significant contribution from the same or neighbouring residue was found in the RdRp + CD10712 system. This finding strengthens the observations of the RMSF result and suggests that the interaction between compound CD10712 and characterized loop was comparatively very poor, due to which binding of compound CD10712 to RdRp has not provided strong support to that loop region, resulting in poor activity.

The binding free energy result has shown that compound DSHS00151 was much more strongly bound to the RdRp with a total binding energy of -107.712 kcal/mol. However, compound CD10712 has shown a reduced binding energy of -91.351 kcal/mol (Figure 5C). Residual decomposition analysis again showed that compound DSHS00151 has a strong non-covalent interaction with residues TRP-800, ILE-802 and HIS-803 (Figure 5). Whereas compound CD10712 was found to be closer to HIS-803 and LYS-805, with an increased decomposition energy for these residues. So, the overall results from the MD simulation study revealed that compound DSHS00151 has a highly stable binding to RdRp in comparison to compound CD10712. The study also showed that both compounds interact with the same characterised loop of the RdRp protein, but compound DSHS00151 has a better interaction, so it stabilized the loop strongly. However, a relatively high deviation was observed in part of the same loop region of the RdRp + CD10712 system with a reduced IC50 value, which strengthened the fact that this characterised loop plays a very essential role in the activity of RdRp.

**Figure 5.**
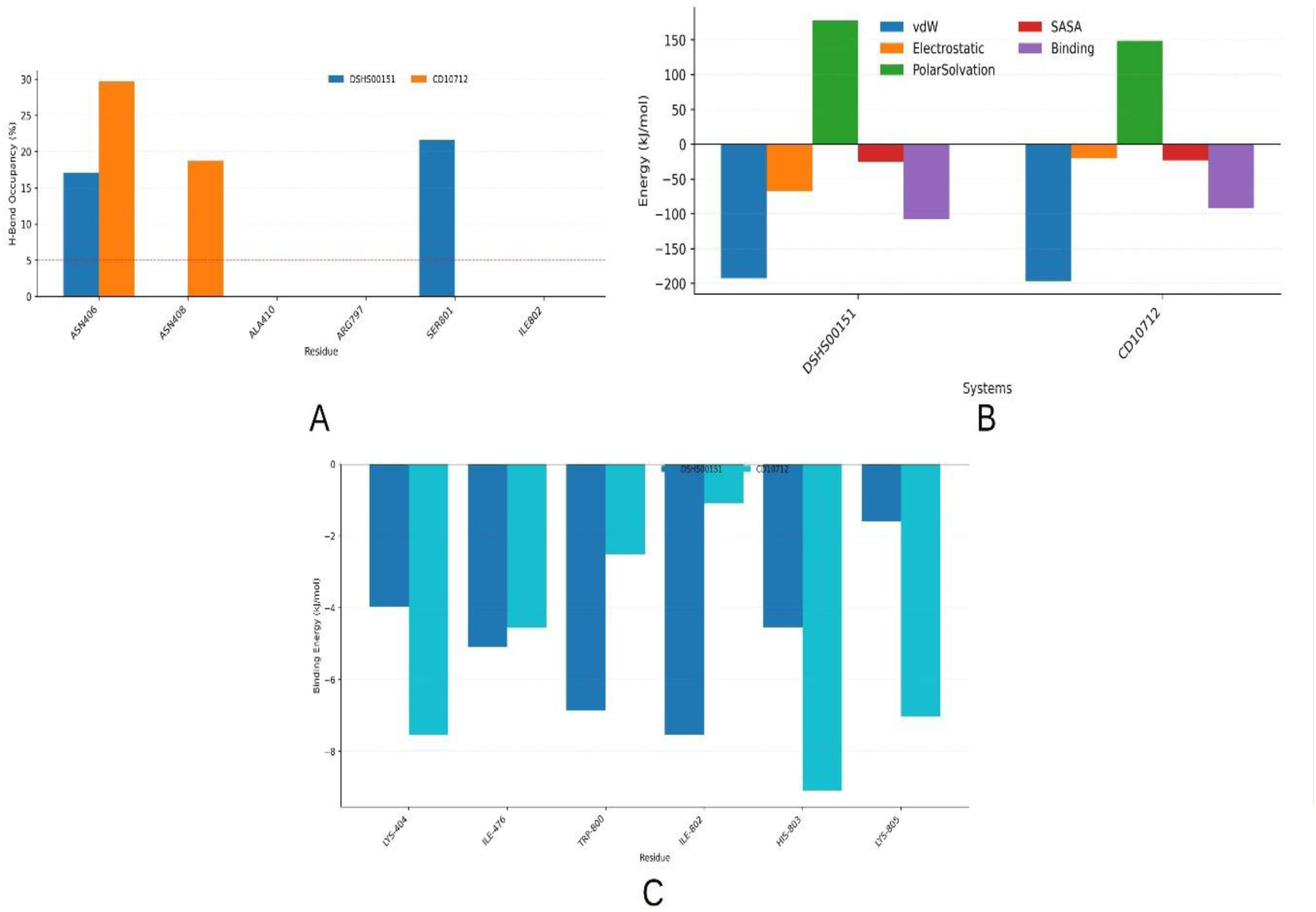
System-wise comparison of (A) significantly contributing residues (contribution of more than 5%) of JEV RdRp in H-Bond formation throughout the simulation time; (B) Different components of binding free energy; and (C) Residual decomposition energy; Systems are shown in different colours, as mentioned in the legend. (For interpretation of the references to colour in this figure legend, the reader is referred to the web version of this article.)

In view of the last conformations from the MD simulation at 100 ns (Figure 6A and B), comparison of the RdRp + DSHS00151 system to the RdRp + CD10712 system has shown a shift in the loop region. (Figure 7) has also been shown that binding of compound DSHS00151 was more inside the pocket, covered by the loop region, whereas binding of compound CD10712 was found more outside, covering the loop region. Comparisons of pre-MD and post-MD ligand conformations (Figure 7) have shown that both compounds have shifted far from the loop region. One end of both compounds was almost at the initial position, while the other end of the molecules was shifted; in the case of compound CD10712, this shift was comparatively high enough to provide flexibility to the loop region, and this may be for the same reason; high fluctuation was observed in the RMSF analysis.

**Figure 6.**
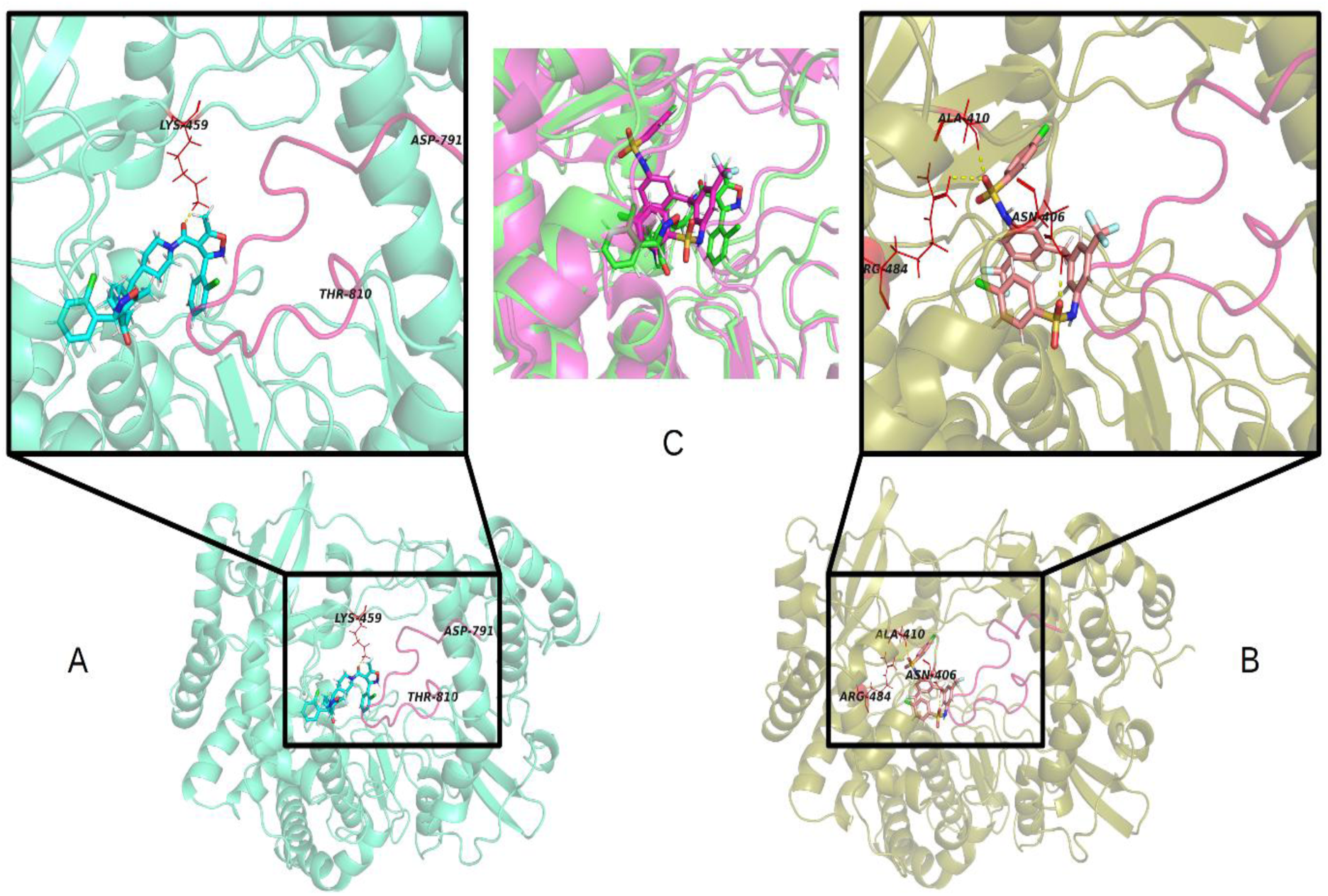
Final conformations (at 100ns) of JEV-RdRp and its interaction pattern with (A) compound DSHS00151 and (B) compound CD10712; along with the (C) comparison of compound DSHS00151 (green) and compound CD10712 (magenta) together at 100ns. (For interpretation of the references to colour in this figure legend, the reader is referred to the web version of this article.)

**Figure 7.**
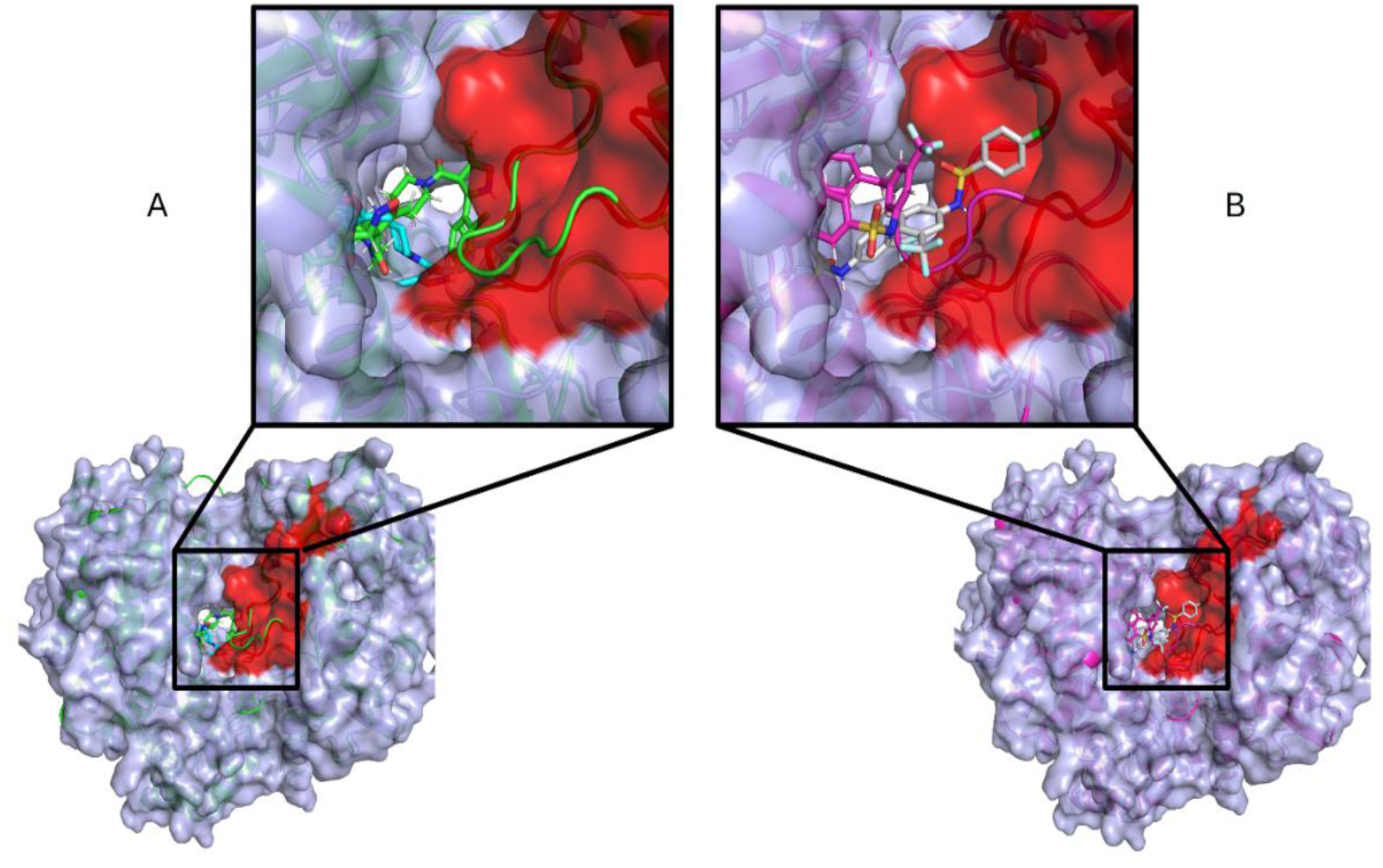
Comparison between the binding of JEV RdRp with (A) compound DSHS00151 at 0th ns (blue) and 100th ns (green) and (B) compound CD10712 at 0th ns (light pink) and 100th ns (magenta). (For interpretation of the references to colour in this figure legend, the reader is referred to the web version of this article.)

The functional significance of the priming loop is further supported by protein–protein interaction analysis involving NS5 and NS3. Previous studies have demonstrated that RdRp activity is regulated through interactions between NS5 and NS3. Consistent with these reports, comparison of apo NS5 with the NS3–NS5 docked complex revealed a pronounced conformational shift in the same priming loop identified in the ligand-binding analyses (Figure 8). The docking model further showed direct contacts between NS3 and the NS5 priming loop, indicating that this region is involved not only in ligand recognition but also in NS3-mediated regulation of polymerase activity. The convergence of ligand-binding, dynamics, energetic, and protein–protein interaction analyses therefore identifies the priming loop as a key regulatory element governing RdRp function. Consequently, compounds capable of strongly engaging and stabilising this loop, such as DSHS00151, are likely to exhibit enhanced antiviral activity and represent promising leads for further optimisation.

**Figure 8.**
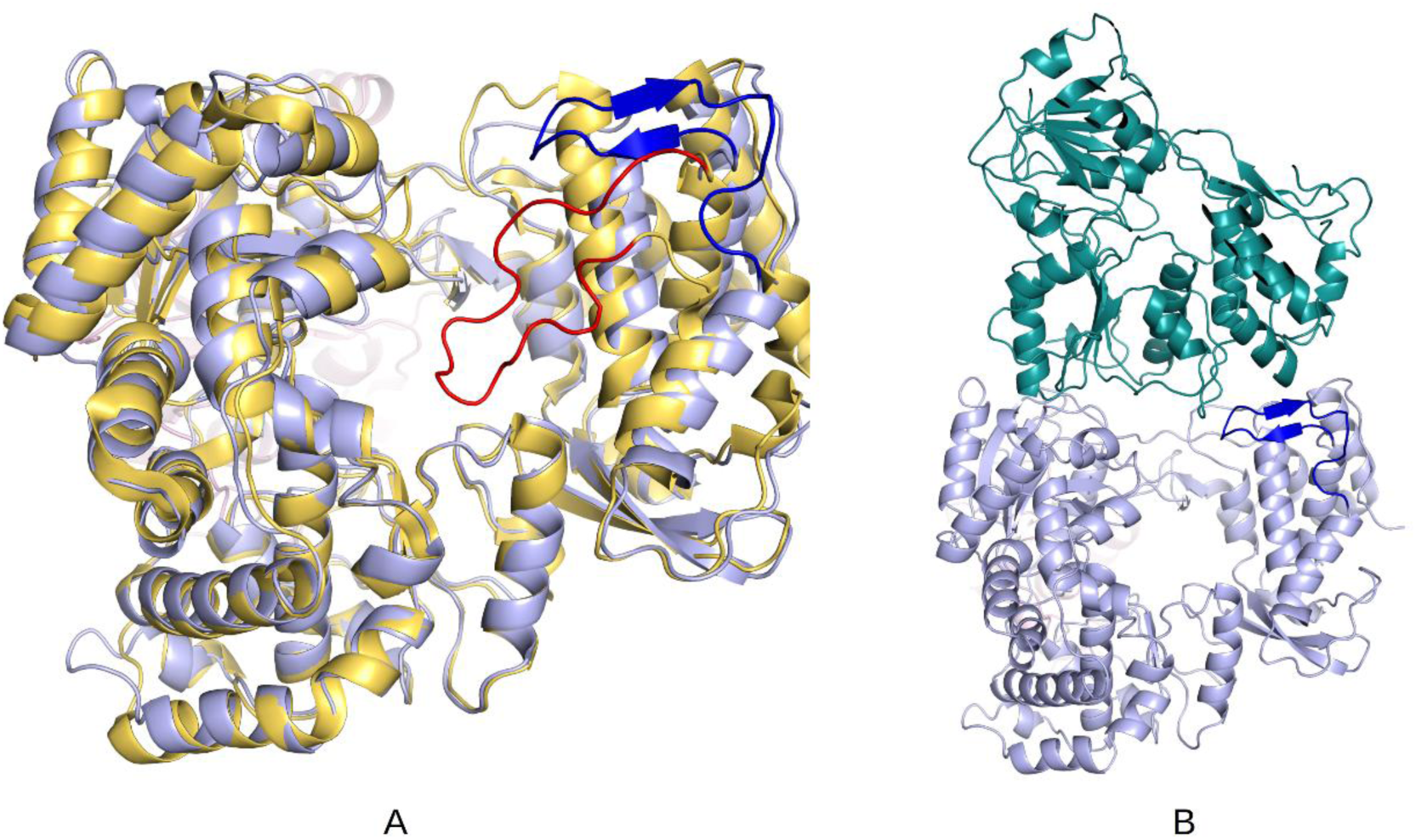
(A) Shift into the JEV-NS5 loop from its APO (red) to JEV-NS3-bound (blue) conformation, along with (B) docked conformation of NS3 (deep teal) and NS5 (blue). (For interpretation of the references to colour in this figure legend, the reader is referred to the web version of this article.)

## Discussion

The RdRp protein in the 3’ domain of NS5 replicates the JEV RNA genome. All flavivirus NS5 RdRps synthesize viral RNA via a “de novo” initiation mechanism, meaning primer-independent replication(Brand et al., 2017). The de novo RNA replication in flavivirus NS5 RdRp is associated with a unique conserved priming loop residue 790-812 inserted in the thumb domain, which regulates RNA-template binding and polymerization, pointing from the thumb subdomain toward the active site (Zhao et al., 2015).The priming loop in NS5 RdRp has been proposed as an allosteric site which takes definitive role in RNA replication(Lim et al., 2016) . Efforts have been made to design inhibitors against the allosteric site, comprising the non-nucleoside reverse transcriptase inhibitor (NNRTI) class of inhibitors affecting viral replication(Celegato et al., 2023; Lim et al., 2016) . Under physiological conditions, the NS5 protein interacts with the NS3 protein to activate RdRp enzyme activity for viral replication(Brand et al., 2024; Xu et al., 2019). The NS5 RdRp priming loop (residues 790-812) function in the replication complex is not well studied.

In the present study, the NS5 crystal structure and protein sequence highlighted in red (residues 790-812) depict a priming loop region, and all JEV genotype NS5 proteins compared show the priming loop to be highly conserved (Figure 1A & B). The NS5 protein ability to interact with NS3 was developed using a mammalian two-hybrid assay, where this interaction could be quantified based on expression of a luciferase reporter gene (Figure 1D). Using a structure-based drug design approach, the inhibitors were screened against the priming loop based on binding energy (Table 1). A total of 15 compounds were screened for NS5-NS3 interaction inhibition and antiviral activity; we found that two compounds, CD10712 and DSHS00151, showed differential interaction inhibition and differential antiviral activity (Tables 2 & 3). The dose-response activity of CD10712 and DSHS00151 showed a direct relationship between priming loop-dependent inhibition of the NS5-NS3 interaction, inhibition of viral infectivity and viral replication in JEV-infected cells (Figure 2 & Figure 3). The varied activity of CD10712 and DSHS00151 on viral replication was due to varied binding affinity of the compounds with the priming loop. Molecular Dynamics (MD) analysis showed that the DSHS00151 compound binding affinity at the priming loop residues is higher than that of the CD10712 compound, suggesting varied antiviral activity (Figure 4). The NS5-NS3 docking studies have predicted that the priming loop comes out from the thumb domain and rotates to bind NS3 for stabilization of the NS5-NS3 complex (Figure 8).

The priming loop residues 790-812 inserted in the thumb domain of NS5 are highly conserved among all JEV genotypes (Figure1B). Since the palm and thumb domains are mostly consistent with the existing WNV and DENV RdRp models, here we focus on the thumb domain of JEV NS5, which has novel observations within. The first reported crystal structure of the full-length JEV NS5 protein shows intra-molecular interactions and key conserved hydrophobic residues at the interface of RdRp and MTase domains to stabilize the unique natural fusion protein, indicating the biological relevance of the observed conformation. The crystal structure also identifies a priming loop residue inserted in the thumb domain, where the thumb is bulkier, carrying additional elements that facilitate de novo initiation (Malet et al., 2007). The priming loop region in NS5 is a single loop structure (residues 790-812) connecting two alpha domains of the thumb for stabilization(Lu & Gong, 2013). The priming loop residues T800 and Arg797 are lined up through cationic-π interaction domains(Brocchieri & Karlin, 1994) in an orientation suitable for priming the initiating NTP. The crystal structure-based information on domain arrangement could not be extrapolated to the NS3 protein interaction complex under physiological conditions. The NS5 protein interaction with NS3 is required to activate the RdRp enzyme for viral replication.

The priming loop function and its involvement in the NS5-NS3 interaction complex are not well studied. In JEV-infected cells, the NS5 and NS3 interaction shown in the replication complex was found to be responsible for viral replication. In other viruses like DENV and ZIKA, the residues at NS5 and NS3 that were involved in interaction and stabilizing the complex(Nannetti et al., 2025; Perera & Kuhn, 2008)Upon mutation of interacting residues, viral replication was inhibited in infected cells(Brand et al., 2024; Tay et al., 2015; Xu et al., 2019). Moreover, the NS5 and NS3 interaction has been determined in yeast two-hybrid assay(Le Breton et al., 2011). Based on the characteristic features of the NS5 and NS3 proteins to interact, we developed a mammalian two-hybrid assay for NS5-NS3 interaction (Figure 1D). This assay demonstrates the physical interaction of NS5 with the NS3 protein in NS5-NS3 expressing HEK-293T cells, which mimics the interaction similar in JEV-infected cells. The functional role of the priming loop in stabilizing NS5-NS3 complex could be studied in a mammalian two-hybrid assay.

The inhibitors identified against the priming loop region for DENV were able to characterize priming loop function in the replication complex required for viral replication. The inhibitors were shown to bind at the active site of the RdRp; this class of compounds were proposed to hinder RdRp conformational changes during its transition from initiation to the elongation process of viral replication (Lim et al., 2016).In our study, using a structure-based drug design approach, we screened Maybridge compounds against the RdRp priming loop pocket residue 790-812. Our high-throughput screening identified 15 compounds based on binding energy that were interacting with the RdRp priming loop (Table 1). Further, all fifteen compounds were screened for NS5-NS3 interaction inhibition in a mammalian two-hybrid assay. Our results showed that two compounds, DSHS00151 and CD10712, showed 2.33 and 1.36-fold reductions in luciferase expression compared to untreated cells in the mammalian two-hybrid assay (Table 2) and showed 0.17- and 0.33-times reduction of infectivity compared to untreated respectively (Table 3). Our studies demonstrate that priming loop involvement in NS5-NS3 interaction stabilization required for functional JEV replication.

Studies show that inhibitors targeting the priming loop can inhibit viral replication in both DENV and Zika (Lim et al., 2016;Tarantino et al., 2016), but no available information regarding NS5-NS3 interaction inhibition.

Dose-dependent analysis establishes the critical relationship between exposure and biological response, guiding compound efficacy. In our studies, dose-dependent analysis of CD10712 and DSHS00151 compounds was done on interaction and antiviral activity. Our results showed that the DSHS00151 compound at 50µM to 1.25µM concentration showed dose-dependent inhibition of NS5-NS3 interaction in a mammalian two-hybrid assay. The interaction inhibition of NS5-NS3 by compound DSHS00151 and CD10712, plotted on a bar graph, showed IC50 3.5µM and 8.701µM respectively (Figure 2A & B). These results suggest that binding of the compound at the priming loop may cause a change in protein conformational structure, resulting in inhibition of interaction. Further, a dose-dependent response for anti-JEV activity of the DSHS00151 and CD10712 compound showed IC50 values of 2.54 µM and 10.45 µM and viral load inhibition of DSHS00151 and CD10712 compound showed IC50 values of 2.9 µM and 18.32 µM respectively. These results validated a direct relationship between the inhibition of RdRp priming loop-mediated interaction with antiviral activity and viral replication inhibition. The varied response of compounds on interaction and antiviral response depends upon the differential binding efficacy of compounds at the NS5 priming loop.

The present molecular dynamics investigation provides mechanistic insights into the differential inhibitory activities of DSHS00151 and CD10712 against JEV RdRp. Although both compounds occupied the same binding pocket, DSHS00151 exhibited superior binding stability, reflected by reduced structural fluctuations, persistent hydrogen-bond interactions and a more favourable binding free energy (Figure 5).

The contribution of residues was investigated through H-bond occupancy results (Figure 5), which showed that residue SER-801, which belongs to the same characterised loop discussed before, was a highly contributing residue for H-bond formation in the RdRp + DSHS00151 system (Figure 5). Ser-801 is highly conserved across all five JEV genotypes. Interestingly, the active compounds targeted against the priming loop of DENV, showing good anti-DENV activity, form H-bonding interactions with His 800 and Glu 802 residues in the priming loop. Amino acid residues of the binding pocket are highly conserved across all DENV serotypes except DENV2. (Yokokawa et al., 2016).

Notably, the enhanced stabilization of the priming loop (residues 791–810) by DSHS00151 suggests that this region plays a central role in maintaining a catalytically competent conformation of RdRp. Previous structural studies have established that the priming loop is essential for de novo RNA synthesis by supporting the positioning of the template and incoming nucleotides during polymerase initiation. Therefore, restricting the conformational dynamics of this loop through ligand binding is likely to interfere with the conformational changes required for efficient RNA synthesis, thereby contributing to polymerase inhibition. The comparatively weaker interactions of CD10712 with the same region may explain its reduced inhibitory potency, highlighting the importance of sustained interactions with the priming loop for effective RdRp inhibition.

The significance of the priming loop is further reinforced by its involvement in NS5–NS3 interactions, which are known to regulate Japanese virus replication. The observed conformational changes in this loop upon NS3 binding (Figure 8) indicate that it functions not only as a structural element within the polymerase active site but also as a regulatory interface during viral replication. The priming loop rotation towards NS3 during interaction implies the stabilization of the priming loop and the NS5-NS3 interaction also. The priming loop-dependent NS5-NS3 interaction, with or without inhibitors, was biochemically studied using our quantitative interaction assay, i.e., the mammalian two-hybrid assay. This result suggests that the priming loop has the property to bind with NS3, as suggested in docking studies. The use of a single residue at the active site was studied by making mutant DENV replicons. The W803A mutant replicon was non-replicative, mutant H800A and Q802A were poorly replicative, and T794A and S796A were more replicative (Lim et al., 2016).

Comparing our results, the DSHS00151 compound interacts with residue S801 of the NS5 priming loop, as identified in MD analysis (Figure 5), and showed inhibition of the NS5-NS3 interaction, resulting in viral replication inhibition. The modulation of the NS5-NS3 interaction in a mammalian two-hybrid assay by high-affinity compounds binding clearly gives an interesting relationship that NS5-NS3 interaction quality activates RdRp enzyme activity for de novo viral replication.

Our study provides convincing evidence that the NS5 RdRp priming loop mediates a critical interaction with NS3 to stabilize the replication complex essential for viral replication. The inhibitors bind to priming loop residues and may cause conformational changes in the NS5 protein, thus inhibiting NS3 interaction, an important mechanism for enzyme function. These findings build on proof-of-concept studies demonstrating the priming loop as a novel allosteric target for developing NNRTI inhibitors and provide a foundation for developing novel JEV therapeutics.

## Author Contributions

**Preeti Mishra:** Writing original draft, Formal analysis, Visualization, Conceptualization. **Shriyanshi Mishra:** Methodology, Investigation, Data Curation, Formal analysis, Writing draft. **Gourav Srivastava:** Writing - review & editing, Methodology, Formal analysis. **Km. Archana:** Review. **Sourav Haldar:** Review & editing. **Mohammad Imran Siddiqi:** Software, Investigation. **Raj Kamal Tripathi:** Writing - review & editing, Supervision, Resources, Investigation, Funding acquisition, Formal analysis, Conceptualization.

## Acknowledgement

Preeti Mishra is thankful to UGC for the Senior Research Fellowship. Shriyanshi Mishra is thankful to the India AI PHD Fellowship. Gourav Srivastava and Km. Archana are thankful to CSIR-CDRI. We express our gratitude to the Repository for Compounds and the Sophisticated Analytical Instrument Facility (SAIF), CSIR-CDRI, for providing Flow Cytometry facilities. We thank Dr Vikas Agrawal at the Sanjay Gandhi Postgraduate Institute of Medical Sciences (SGPGIMS), Lucknow, for providing the Japanese Encephalitis Virus. Raj Kamal Tripathi acknowledges funding from CSIR-CDRI (IHP0022). We acknowledge the use of AI for grammar correction.

